# Broad Sarbecovirus Neutralization by an S2-Directed Plasma Antibody Defines a New Site of Vulnerability in SARS-like Viruses

**DOI:** 10.64898/2026.06.06.730562

**Authors:** Douglas R. Townsend, Ling Zhou, Michael L. Mallory, William N. Voss, Sean A. Knudson, Cory Acreman, John M. Powers, Miranda L. Hubbard, Alexandra Schafer, Nick Catanzaro, David R. Martinez, Sumit Pareek, Connor Mullins, Annalee W. Nguyen, Jennifer A. Maynard, Ralph S. Baric, George Georgiou, Jason S. McLellan, Gregory C. Ippolito, Jason J. Lavinder

**Author notes:** Authors contributed equally (Douglas R. Townsend, Ling Zhou, Michael L. Mallory).

## Abstract

The repeated emergence of pathogenic coronaviruses highlights the continual threat of zoonotic spillover events and the need to understand conserved regions of the viral spike protein that influence immune recognition. Here, we employ high-resolution proteomic analysis of circulating immunoglobulins to characterize the antibody response to the SARS-CoV-2 spike protein in vaccinated and infected individuals. We recombinantly expressed abundant plasma IgG lineages and identified nine antibodies targeting the highly conserved S2 spike subunit. Most of these S2-reactive plasma mAbs (7 of 9) exhibited neutralizing activity against pangolin coronavirus, while one mAb (SC45) demonstrated potent (IC_50_ < 1 µg/mL) neutralizing activity against both pangolin and bat sarbecoviruses RsSHC014 and WIV1. Additionally, SC45 exhibited potent prophylactic efficacy in the K18-hACE2 mouse model challenged with Pangolin-Guangdong CoV MP789 (Pg-CoV), significantly reducing lung viral replication at day 4 post-infection. The cryo-EM structure of SC45 bound to the prefusion-stabilized S2 domain of Pg-CoV reveals that SC45 targets a novel, conserved epitope in the connector domain. Strikingly, even though SC45 was the most abundant antibody (∼13%) of anti-spike plasma IgG in an infected patient and binds to both SARS-CoV-2 and Pg-CoV spike with comparable affinities, it fails to neutralize early (D614G and Omicron BA.1) SARS-CoV-2 variants. Together, these findings identify a conserved site of vulnerability in the spike S2 subunit which informs on potential mechanisms of antibody immunity to zoonotic coronaviruses.

## Introduction

The repeated emergence of pathogenic coronaviruses such as SARS-CoV-1 (2002), MERS-CoV (2012), and SARS-CoV-2 (2019) underscores the persistent threat of zoonotic coronavirus (CoV) spillover and the need for broadly protective countermeasures^1–5^. Both SARS-CoV-1 and SARS-CoV-2 belong to the betacoronavirus subgenus *Sarbecovirus*, which can be classified into four evolutionary clades based on their spike glycoprotein receptor binding domain (RBD) sequence^6,7^. Clades 1a and 1b include multiple “SARS-like” strains considered high-risk for zoonotic spillover and pandemic potential due to their ability to engage human angiotensin-converting enzyme 2 (hACE2) for cell entry and their efficient replication in primary human airway epithelial cells^8–11^. The rich reservoir of SARS-like viruses with documented human-cell infection capacity, coupled with their high recombination rates and frequent interface with human populations highlights the need for the development of broadly protective countermeasures to blunt future SARS-like outbreaks. To effectively develop broadly protective and durable vaccines and therapeutics against diverse SARS-like viruses, it is critical to understand conserved epitopes between viral strains, their cross-reactive breadth, and protective capacities.

The CoV spike glycoprotein mediates receptor binding and membrane fusion and is the principal target of neutralizing antibodies^12–15^. The spike protein comprises an S1 subunit, responsible for host receptor recognition, and an S2 subunit which encodes the highly conserved fusion machinery required for viral entry^12,16^. Although neutralizing antibody responses to the spike protein primarily target the S1 domain, the S2 subunit exhibits less sequence variability across CoVs and thus represents a promising target for vaccine design to elicit broad-CoV immunity^17^.

Within the S2 subunit, three major protective antibody epitopes have been described: the stem-helix, the fusion peptide, and the S2-apex. Stem-helix antibodies recognize a conserved helical bundle near the base of the spike, neutralize SARS-CoV-2 in vitro with IC50s in the low-to-mid µg mL⁻¹ range^17–20^ and provide protection in animal models at least in part through Fc-mediated effector functions^17,18,21,22^. Fusion-peptide antibodies bind a conserved region adjacent to the S2′ cleavage site and can neutralize both alpha- and betacoronaviruses, although with lower potency than stem-helix directed antibodies. Similar to stem-helix antibodies, fusion peptide-directed antibodies also confer robust prophylactic protection against SARS-CoV-2 in mice and hamsters^23,24^. Antibodies directed to the S2-apex generally lack measurable neutralizing activity but protect against SARS-CoV-2 challenge *in vivo* through Fc-dependent mechanisms^25,26^.

S2-directed antibodies have been shown to be highly abundant within the polyclonal IgG response in SARS-CoV-2 convalescent individuals^27,28^. Despite their comparatively modest neutralization potencies, depletion of S2 antibodies from plasma has been shown to result in reduced protection ^23,29,30^. Additionally, pre-existing memory B cells primed by infection with betacoronaviruses have been shown to be activated upon infection with SARS-CoV2 and to encode cross-reactive non-neutralizing S2-specific antibodies capable of mediating protection via Fc-effector functions in mouse challenge experiments^31–33^.

Here, we report the discovery of a circulating human antibody that targets a novel site of vulnerability in the spike protein S2 subunit of sarbecoviruses. Using liquid chromatography-tandem mass spectrometry (LC-MS/MS) serum antibody proteomics (Ig-Seq) ^27,34–37^ on a cohort of SARS-CoV-2 infected and vaccinated donors, we identified a highly abundant S2-directed plasma antibody, SC45, which neutralizes members of both clade 1a and clade 1b sarbecoviruses, including SARS-CoV as well as several high-risk zoonotic strains. We show that SC45 confers *in vivo* prophylactic protection against infection with Pangolin-Guangdong CoV (Pg-CoV) and that even though it did not neutralize SARS-CoV-2 Wuhan-Hu-1 it did confer partial protection when administered prophylactically, presumably via Fc functions. Structural characterization revealed that SC45 binds to a conserved hairpin loop in the connector domain (CD) of sarbecovirus spike glycoproteins. Together, these results reveal a novel site of vulnerability within the CD of SARS-like CoVs.

## Results

### Isolation and characterization of circulating S2-binding antibody lineages

We characterized the plasma antibody repertoire targeting the SARS-CoV-2 spike protein (stabilized SARS-CoV-2 via the introduction of six Pro substitutions, hereinafter referred to as S6P) in four SARS-CoV-2 vaccinated and infected subjects^27^ using the Ig-Seq LC-MS/MS proteomics pipeline^38^ (**Figure 1A** and **Figure S1**). Ig-Seq provides a readout of the identities and relative abundances of antibodies that comprise the polyclonal plasma response to an antigen of interest. Two of these subjects experienced subsequent secondary breakthrough infection. Analysis of serological persistence of distinct antibody lineages across three successive antigenic exposures indicates that the circulating anti–SARS-CoV-2 spike IgG repertoire undergoes extensive reshaping, such that only a subset of IgG lineages persist across exposures, while newly detected plasma IgG lineages (arising either from memory B-cell recall or from *de novo* elicitation) constitute a substantial fraction of the response. In donor P25 (**Figure 1B**), 15 lineages comprising 33% of the breakthrough 1 plasma IgG repertoire were initially observed in the post-vaccination plasma IgG response. This donor subsequently experienced a second breakthrough infection (breakthrough 2), in which 10 lineages accounting for 23% of the breakthrough 2 IgG repertoire were derived from lineages initially detected in breakthrough 1 infection plasma. Four lineages accounting for 9% of the breakthrough 2 repertoire were detectable across all exposure events. In donor P33 (**Figure 1C**), 8 lineages accounting for 28% of the breakthrough 1 plasma response were initially observed in the post-vaccination repertoire. Together, these patterns indicate substantial lineage turnover in the polyclonal IgG against SARS-CoV-2 spike with only a limited subset of lineages retained across all antigenic exposures.

**Figure 1:**
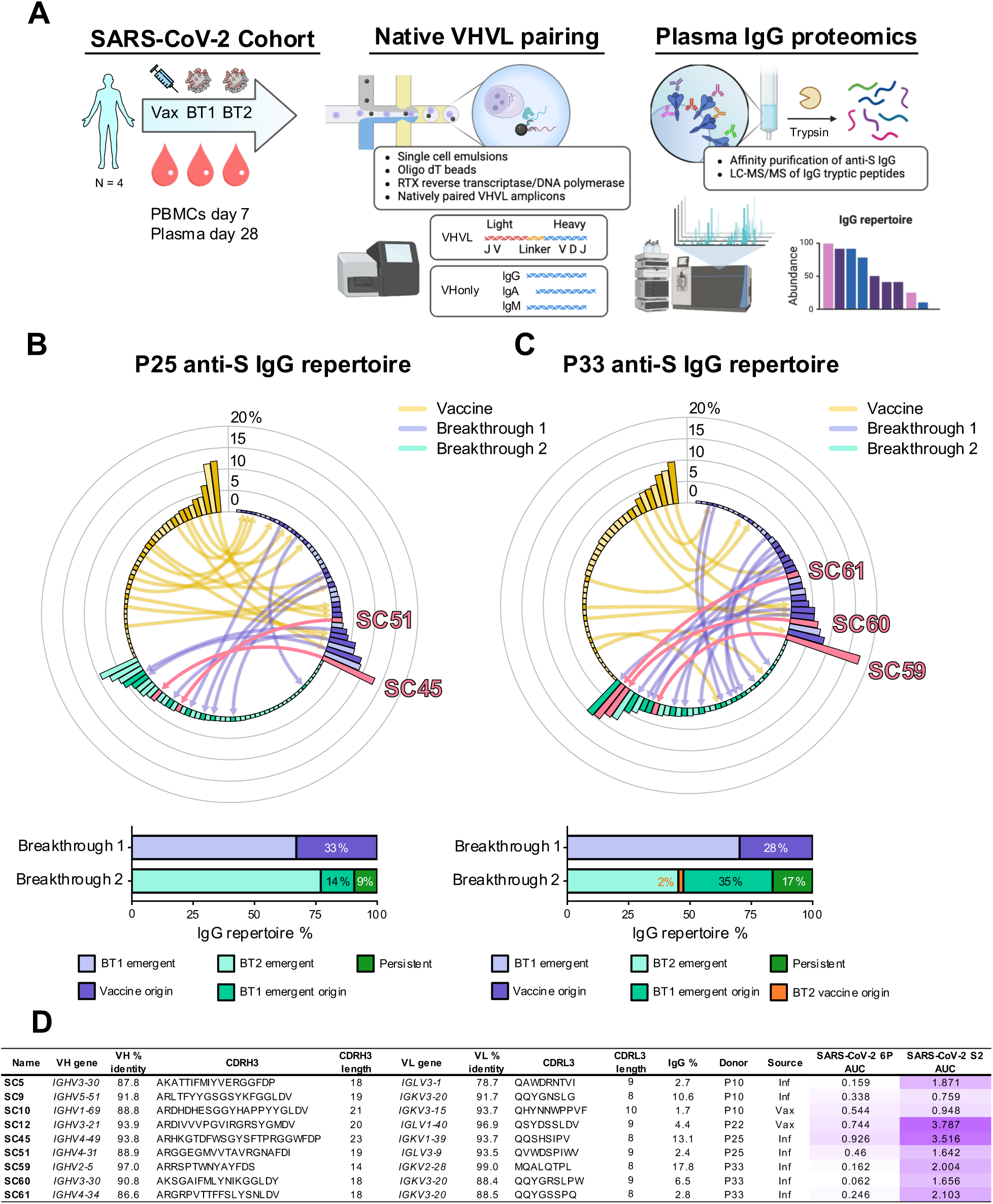
Profiling longitudinal IgG serological repertoires across SARS-CoV-2 exposures identifies abundant S2-directed antibodies. (A) Ig-seq analysis pipeline of anti-SARS-CoV-2 spike IgG following vaccination and breakthrough infections in donors P25 and P33. Image created with BioRender.com (B-C) Serological repertoires of SARS-CoV-2 spike-specific IgG in plasma in donors P25 (left) and P33 (right) across SARS-CoV-2 exposures. Each bar denotes an individual antibody lineage plotted by its relative abundance, showing only lineages that account for >0.5% of the spike-specific IgG repertoire. Directional arrows connecting bars in the histogram demonstrate recalled lineages at each time point. The proportion of plasma IgG which account for overlap between time points are shown below and shaded as follows: vaccine and BT1 (dark purple), vaccine and BT2 (orange), new lineages emerged in BT1 overlapping with BT2 (teal), and persistent across all exposures (green). (D) Immunogenetic features of a panel of S2-directed antibodies identified by Ig-seq (rows) and ELISA AUC of antibodies against SARS-CoV-2 6P and SARS-CoV-2 S2 antigens.

We recombinantly expressed and screened n=60 plasma-derived mAbs as IgG1 from these four donors by indirect ELISA against full length prefusion stabilized S6P. As full-length SARS-CoV-2 spike transiently exposes cryptic S2 epitopes and several S2 directed antibodies exhibit reduced binding to S6P, we also screened the plasma-derived mAbs against SARS-CoV-2 spike Hexapro-SS, a prefusion-stabilized S2-only antigen engineered to lock the S2 subunit in a prefusion trimer via six proline mutations, interprotomer disulfide bonds and an engineered salt bridge (Hexapro+S704C/K790C+Q957E) hereafter referred to as S2-antigen^39^ (**Figure 1D**). We found n=9 S2-specific antibodies which exhibited stronger binding to the SARS-CoV-2 S2-antigen as compared to S6P (**Figure 1D**). These S2-specific mAbs were tested for live-virus neutralization against a panel of clade 1a and 1b sarbecoviruses that are considered pre-emergent zoonotic risks as well as distantly related MERS-CoV and HCoV-OC43 betacoronaviruses (**Figure 2A**). While none of the nine S2-directed mAbs neutralized SARS-CoV-2 D614G or Omicron BA.1, 6 of 9 (SC5, SC9, SC45, SC59, SC60, SC61) were able to neutralize SARS-CoV-1, and a subset (SC9, SC45, SC59, SC60, and SC61) also neutralized Pangolin Guangdong^8^ (Pg-CoV), a clade 1b zoonotic virus (**Figure 2B** and **Figure S2**). Overall, seven of the nine S2-directed mAbs exhibited neutralizing activity against at least one pre-emergent zoonotic virus (clade 1a and 1b sarbecoviruses which circulate in wildlife reservoirs), with Pg-CoV demonstrating the greatest sensitivity to antibody-mediated neutralization among our panel of S2 mAbs. None of the S2 mAbs tested here neutralized MERS nor HCoV-OC43 suggesting that their neutralization profile is restricted to sarbecoviruses.

**Figure 2:**
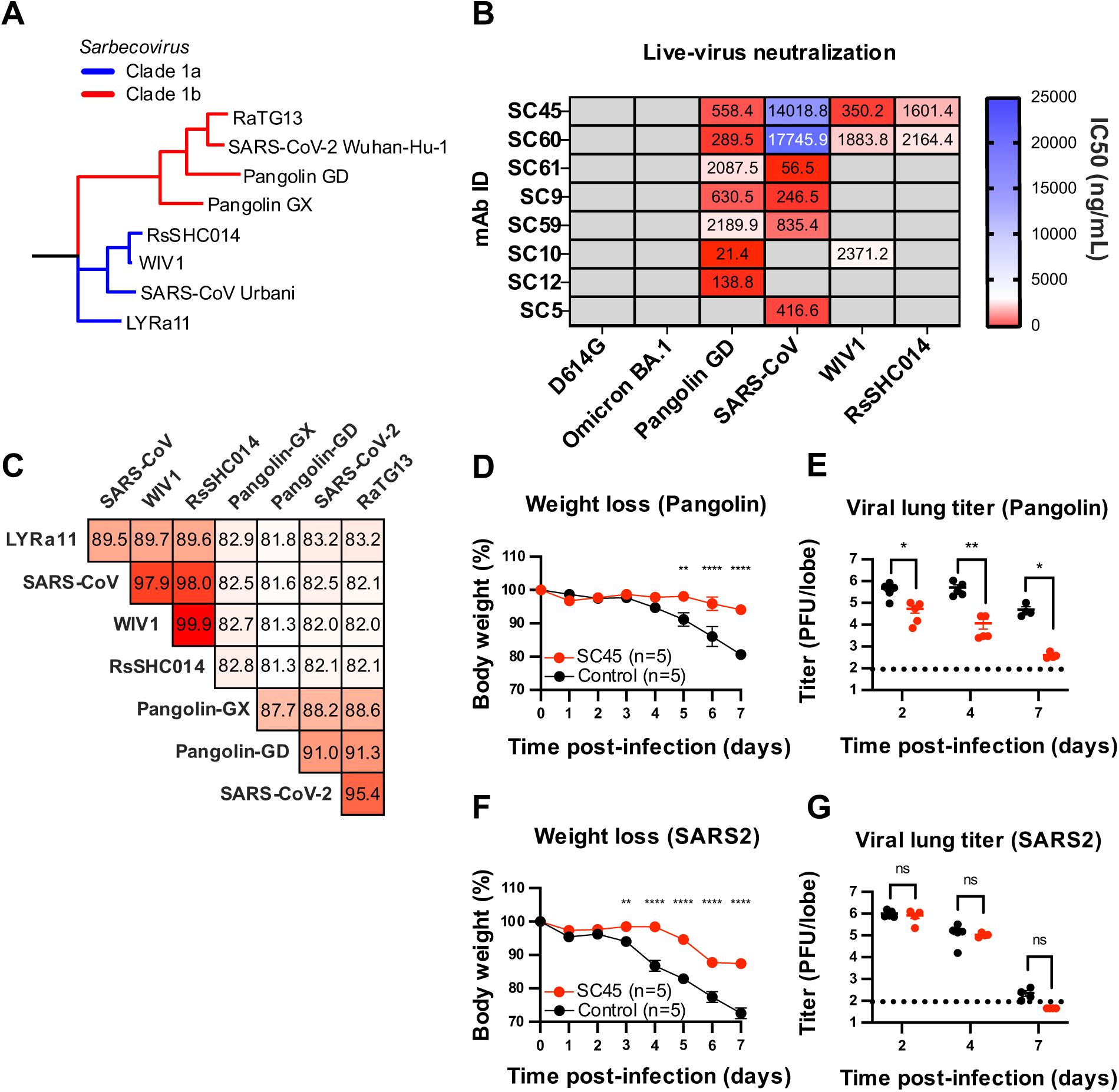
SC45 is a potent and broadly neutralizing S2-directed antibody and protects against disease in K18-hACE2 mouse models of infection. (A) phylogenetic tree of spike sequences from select members of the Sarbecovirus subgenus (Homo sapiens: SARS-CoV Urbani, SARS-CoV-2 Wuhan-Hu-1; Rhinolophus sinicus: WIV1, RsSHC014; Rhinolophus affinis: LYRa11; Manis javanica: Pangolin Guangdong, Pangolin Guangxi/P2V). Members of clades 1a (red) and 1b (blue) are indicated by color; blu. (B) Live-virus neutralization assays using the S2-directed antibody panel. Non-neutralizing activity is shown in solid gray (Fig. 1D) (C) Sequence identity matrix for the S2-subunit of clade 1a and 1b sarbecoviruses. (D-G) In vivo prophylactic protection of K18-hACE2 mice challenged with 10E4 plaque-forming units (PFUs) of Pangolin-Guangdong CoV MP789 or SARS-CoV-2 Wuhan-Hu-1 treated with 200 ug (8 mg/kg) of SC45 or isotype control antibody DENV2-2D22. One mouse from the Pg-CoV treatment group died from anesthesia-related complications on day 4 and one mouse from the Pg-CoV control group died from infection on day 6. Weight change of mice over time represented as mean +/− S.E.M. for n = 15 day 1-2, n = 10 day 3-4, n = 4 (Pg-CoV) and n=5 (SARS-CoV-2) day 5-7. Comparison of percent weight loss with the isotype control was performed using a two-way analysis of variance (ANOVA) followed by Sidak’s multiple comparison test: * P<0.05, ** P < 0.01, *** P < 0.001, **** P < 0.0001. Comparison of lung viral burden was assessed by plaque assay. Data consists of the mean +/− S.E.M and significant differences calculated using the Mann-Whitney U test: * P < 0.05, ** P < 0.01. All neutralization assays were run in duplicate.

The broadest neutralization was observed for SC45 and SC60, which potently neutralized clade 1b Pg-CoV and multiple clade 1a viruses (WIV1, RsSHC014), while exhibiting weaker neutralizing activity against clade 1a SARS-CoV-1. While Pg-CoV and SARS-CoV-2 share 91% amino acid identity in the S2 domain, SC45 and SC60 neutralize the former, but not the latter (**Figure 2C**). However, both antibodies neutralized the more divergent clade 1a viruses (82-83% amino acid S2 sequence identity).

### SC45 Confers *in vivo* Protection Against Pangolin-CoV Infection and Partial Protection Against SARS-CoV-2

We next sought to explore the protective efficacy of SC45 as a prophylactic in the transgenic K18-hACE2 mouse models of Pg-CoV and SARS-CoV-2 infection. Three groups of n=5 mice per group were administered either 200 ug (8mg/kg) of mAb DENV2-2D22 (isotype control; anti-dengue serotype 2 envelope protein) or SC45. Mice were challenged intranasally 12 hours after antibody infusion and monitored daily for weight loss. Animals were sacrificed at day 2, day 4 or day 7 post-challenge, and lung tissues were harvested to determine viral lung titers. One mouse in the control group was found dead in its cage from infection on day 6, and one mouse in the treatment group died of anesthesia-related complications on day 4. These two mice were excluded from the final longitudinal weight loss and day 7 viral titer analyses. Compared to the isotype control antibody DENV2-2D22, SC45-treated Pg-CoV infected mice showed substantially reduced weight loss (5.9% vs 19.4%) at day 7 suggesting a protective role for SC45 (**Figure 2D**). Consistent with the reduced weight loss, viral lung titers were substantially lower in SC45 treated mice compared to the isotype control (**Figure 2E**). Despite its lack of neutralization activity against SARS-CoV-2 *in vitro*, SC45 also conferred partial protection against SARS-CoV-2 challenge *in vivo*, attenuating weight loss (12.6% vs 27.5%) compared to the isotype control at day 7 (**Figure 2F**), but had no effect on lung viral titers (**Figure 2G**).

### SC45 Binding is Reduced by Prefusion Stabilizing Spike Mutations

To examine the binding properties of SC45 to the S2 domain, we performed surface plasmon resonance (SPR) using the SARS-CoV-2 S2-antigen^39^. SC45, as a monovalent Fab, bound to SARS-CoV-2 S2-antigen with nanomolar affinity (K_D_ = 58.2 nM) (**Figure 3A**). As the SARS-CoV-2 full length spike has been shown to transiently expose cryptic S2 epitopes^39^, we sought to determine whether binding differs between full length S6P spike and S2-antigen. SC45 bound to S6P with substantially lower affinity (K_D_ = 1.4 uM) (**Figure 3B**). Evaluation of SC45 binding to a SARS-CoV-2 S6P construct engineered to maintain a fully closed “three RBD-down” conformation via the introduction of interprotomer disulfide bonds (S6P-3D; residues) revealed no detectable binding (**Figure 3C**). We next adapted the SARS-CoV-2 S2-antigen and S6P sets of prefusion-stabilizing mutations to Pg-CoV to generate Pg-CoV S2-antigen and Pg-CoV S6P (proline mutations at residues 809, 884, 891, 934, 978 and 979). Analysis of SC45 bound to Pg-CoV S2-antigen and Pg-CoV S6P by SPR confirmed a similar trend observed in the SARS-CoV-2 constructs, wherein SC45 binds to the S2-antigen with substantially higher affinity than full-length prefusion stabilized Pg-CoV S6P (K_D_ = 28.7 nM vs 303.5 nM) (**Figure 3D-E**). To determine whether the S2 binding properties of SC45 are consistent across a more distant clade 1a sarbecovirus, we adapted the S2 stabilizing strategy to WIV1 to produce WIV1 S2-antigen. SC45 bound to WIV1 S2-antigen with comparable affinity to Pg-CoV and SARS-CoV-2 S2-antigens (K_D_ = 32.1 nM) (**Figure 3F**).

**Figure 3:**
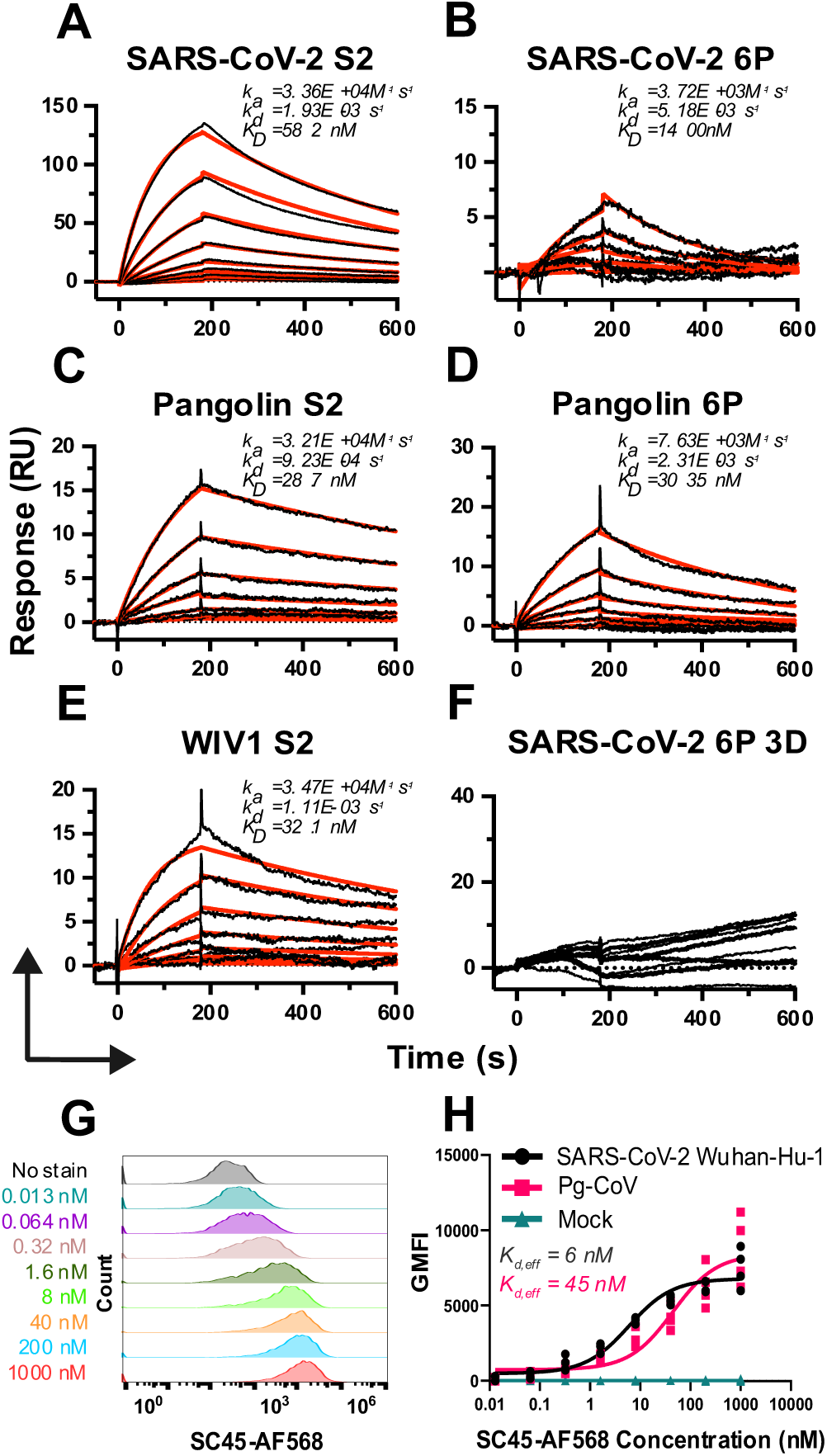
SC45 exhibits reduced binding affinity for full-length prefusion stabilized spike antigen. (A-F) SPR sensorgrams of SC45 Fab binding to prefusion stabilized S2 antigen of SARS-CoV-2, Pg-CoV, WIV1 and prefusion stabilized full length spike antigens of SARS-CoV-2 and Pg-CoV. (G) Expi293 cells were transiently transfected with plasmids encoding EGFP and either SARS-CoV-2 Wuhan-Hu-1 or Pg-CoV then incubated with 0-1000 nM of SC45-AF568 before measurement of GMFI by flow cyometry. (H) GMFI data fit a three-parameter logistic curve to determine the effective K_d_ (K_d,eff_) for SARS-CoV-2 Wuhan-Hu-1 (black), Pg-CoV (pink) and mock EFGP-only transfected cells (teal).

The SPR data indicate that the SC45 epitope may be partially occluded in the context of full-length spike or that a greater degree of spike stabilization decreases binding affinity, consistent with reports that prefusion stabilizing mutations can reduce the accessibility of S2 epitopes^23,24,40,41^. To evaluate binding to authentic, wild-type S protein lacking any stabilizing mutations, we transiently expressed SARS-CoV-2 or Pg-CoV S protein on the surface of Expi293 cells (**Figure S3**). Surface-expressed spike is expected to more accurately recapitulate *in vivo* antibody interactions with the authentic antigen encountered during natural infection. Cells were incubated with 0.06 – 1000 nM of AF568-labeled SC45 antibody (**Figure 3G**), and the mean fluorescence intensity (MFI) was measured by flow cytometry to determine a steady state effective *K*_D_ ^25,42^. SC45 demonstrated robust binding to wt SARS-CoV-2 spike protein (effective *K*_D_ = 6 nM) and somewhat lower binding to wt Pg-CoV (effective *K*_D_ = 45 nM) (**Figure 3H**). We conclude that SC45 binds well to wt S protein in a manner comparable to the stabilized S2-antigen but further stabilization, as achieved in the S6P and S6P-3D constructs, leads to a reduction in binding affinity.

### SC45 binds to a novel epitope in the spike protein connector domain

To define the molecular basis of SC45 binding and neutralization, we solved the cryo-EM structure of SC45 Fab in complex with Pg-CoV S2-antigen stabilized construct at 2.91Å. We solved the structure with prefusion-stabilized Pg-CoV S2 since SC45 neutralizes the pangolin live virus with high potency unlike the SARS-CoV-2 live virus.

When in complex with SC45, Pg-CoV S2-antigen unexpectedly adopted a postfusion-like conformation, even though it contains prefusion-stabilizing mutations (**Figure 4A**)^43^. The SC45 epitope is located within the connector domain (CD) of the S2 subunit (residues 1076-1114 SARS-CoV-2; 1068-1106 Pg-CoV). In the prefusion state, the CD is situated at the most basal part of the spike, proximal to the viral membrane, immediately downstream of HR1 and preceding the stem-helix and HR2 (**Figure 4A**). During fusogenic rearrangements the CD serves as a structural hinge, facilitating the reorientation of HR2 as it zippers along HR1, ultimately driving membrane fusion^44^.

**Figure 4:**
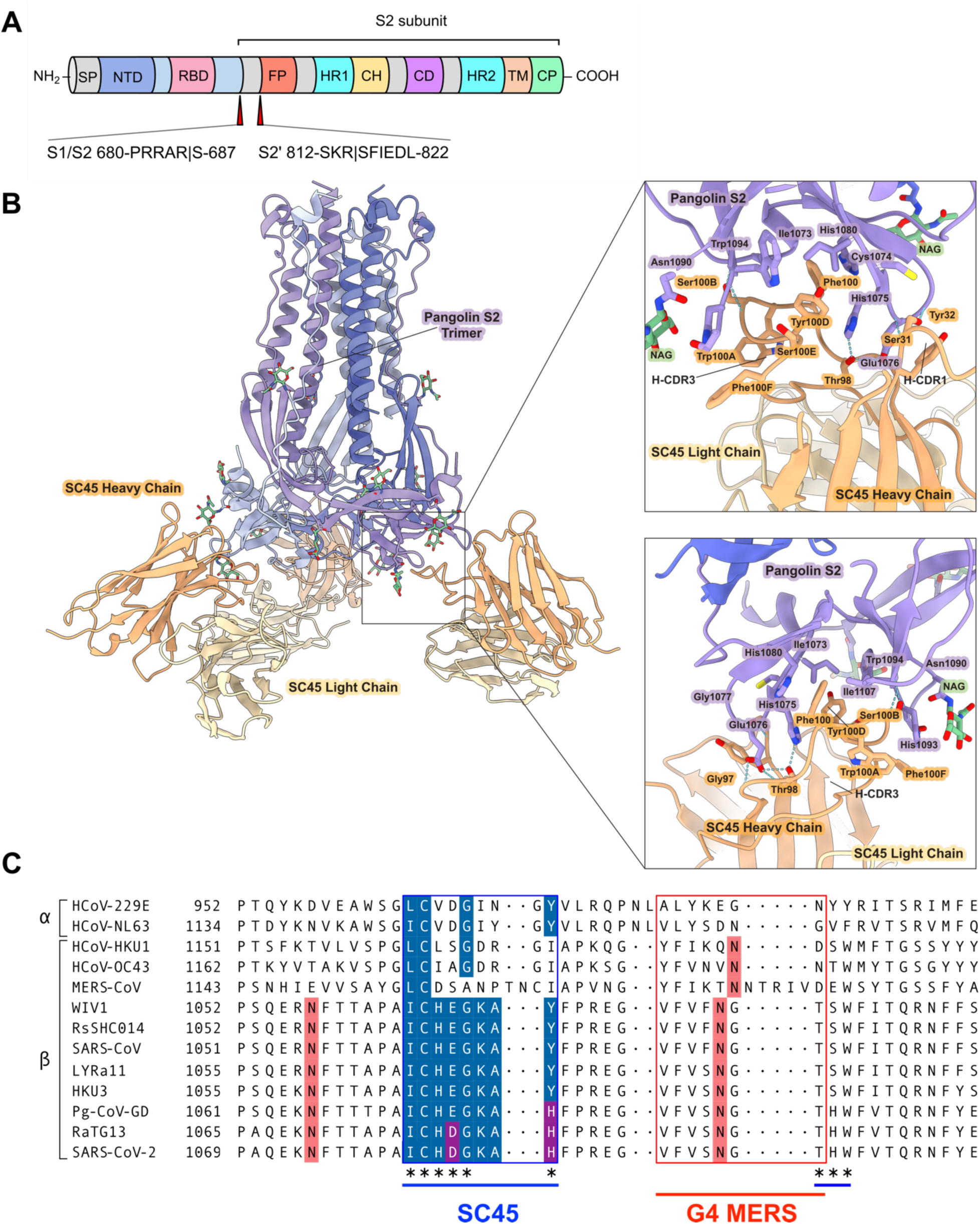
Cryo-Em structure of the SC45:Pg-CoV S2-antigen complex. (A) Schematic of CoV spike protein domain organization with the S1/S2 and S2’ cleavage sites for SARS-CoV-2 labeled. (B) Cryo-EM structure of the SC45 Fab bound to the postfusion conformation of the Pg-CoV S2 subunit (purple). The CDRH3 of SC45 inserts into a hydrophobic pocket, with key residues shown as sticks; Phe100 makes extensive hydrophobic contacts within the pocket. Glu1076 is highlighted as the sole epitope residue that differs between Pg-CoV and SARS-CoV-2. (C) Multiple sequence alignment of selected alpha- and beta-coronaviruses. SC45 and MERS G4 epitope regions are indicated by blue and red boundaries, respectively. Conserved residues within the SC45 epitope are shown in blue, with conserved substitutions in purple. Asterisks denote residues forming significant contacts with the SC45 Fab in the Pg-CoV structure. N-linked glycosylation sites are shown in salmon.

SC45 engages the Pg-CoV S2-antigen with a binding mode dominated by the heavy chain, while the light chain does not participate in the binding interface. The antibody buries a total surface area of 602 A^2^ with interactions primarily mediated by residues derived from a germline IGHD3-3 gene segment within CDRH3. Of note, Phe100 (Kabat numbering) and Tyr100D within the IGHD3-3 segment motif -DFWSY-inserts into a deep hydrophobic pocket formed by Ile1073, His1075, His1080. This interaction is further stabilized by predicted π–π stacking interactions between Phe100 of CDRH3 and His1080 of S2 (**Figure 4B**). This hydrophobic “hook” is flanked by Thr98, which forms polar interactions with Glu1076, further anchoring the CDRH3 loop. CDRH2 contributes additional contacts through Ser31 and Tyr32, which engage a linear segment spanning residues Cys1074 to Gly1077 which is largely conserved across sarbecoviruses (**Figure 4C**). Within the Pg-CoV spike, Cys1074 forms a disulfide bond with Cys1118, which is well conserved across beta and alpha coronaviruses. Interestingly, the SC45 light chain does not make any contacts with the post-fusion S2 conformation in our structure. This heavy chain dominant mode of recognition is reminiscent of the CD-binding mouse antibody G4, which neutralizes MERS-CoV through heavy-chain dominated recognition of the CD^45^ (**Figure 4C**).

To investigate SC45’s apparent dual prefusion and postfusion binding, we aligned the primary contacting residues within our complex to those of the prefusion Pg-CoV structure (PDBID: 7BBH). This revealed that the SC45 betahairpin epitope structure is retained between prefusion and postfusion states despite substantial displacement of the CD region during fusogenic rearrangements (**Figure S4**). Although SC45 exhibits high affinity for the full-length wt Pg-CoV spike, it is observed bound to S2 in a postfusion-like conformation by cryo-EM. Together, these findings support a model in which SC45 engages the metastable S2 domain, funneling intermediate states towards postfusion-like rearrangements.

There is a high degree of sequence conservation in the SC45 binding epitope among sarbecoviruses (**Figure 4C**) with 6 of 9 key SC45 contact residues being identical across clade 1a and clade 1b sarbecoviruses. In our structure the SC45 Fab is flanked by glycosylation sites at N1090 (one from an adjacent promoter) which SC45 avoids via its angle of approach (**Figure 4B**). Two notable mutations were identified within contact residues of the SC45 epitope in SARS-CoV-2 relative to Pg-CoV: E1084D and Y1088H, with the E1084D substitution distinguishing the clade 1a viruses SARS-CoV-2 and RaTG13 from Pg-CoV. To determine if this conservative substitution was responsible for the lack of SC45 neutralization, a SARS-CoV-2 mutant was engineered containing the D1084E mutation. This single amino acid substitution effectively converts the SC45 epitope to the corresponding Pg-CoV sequence. However, SARS-CoV-2 D1084E was also not neutralized by SC45 (**Figure S5**). Therefore, other features of the S2 domain must dictate susceptibility to SC45-mediated neutralization of other tested sarbecoviruses.

It should be noted that SARS-CoV-2 and Pg-CoV also differ in their respective entry pathways^46–49^ since SARS-CoV-2, but not Pg-CoV, possesses a polybasic furin cleavage site at the S1/S2 junction^50,51^ which enables pre-processing during viral packaging and TMPRSS2 activation at the plasma membrane^52^. Pangolin and bat sarbecoviruses (Pg-CoV, RsSHC014, WIV1) lack the furin cleavage site and are reported to rely on endosomal cathepsin for fusion activation^53–56^. Because these differences in proteolytic processing of spike alter the timing and location of membrane fusion, we hypothesized that SC45-mediated neutralization may depend upon the viral endosomal entry pathway and that redirecting SARS-CoV-2 to utilize endosomal entry could render it susceptible to SC45. To test this, we performed neutralization assays using a SARS-CoV-2 furin knockout variant (ΔPRRA), which has been shown to redirect viral particles to the endo/lysosomal entry pathway in Vero cells^51^. Nonetheless, SC45 failed to neutralize the furin knockout SARS-CoV-2 virus (**Figure S5**). Collectively, these results indicate that neither epitope substitution nor forcing SARS-CoV-2 to enter through the endosomal pathway sensitize SARS-CoV-2 to SC45-mediated neutralization. Given the comparable binding of SC45 to wt Pg-CoV and wt SARS-CoV-2 Wuhan-Hu-1 (**Figure 3**), these data suggest SC45 may neutralize through disruption of a long-range allosteric network coupled to the fusion process, which may differ across sarbecoviruses^57^.

## Discussion

Our study identifies a previously unrecognized site of vulnerability within the sarbecovirus S2 connector domain. While the hinge-like function of the connector domain during rearrangements from prefusion to postfusion is a conserved feature of all coronavirus spike proteins, this region adopts distinct structural conformations across CoV subgenera and the specific epitope we identified is highly conserved within sarbecoviruses. We show that SARS-CoV-2 vaccination and breakthrough infection can elicit highly abundant spike S2-specific plasma IgG lineages. Interestingly, although abundant within the plasma, none of these S2-specific IgG were able to neutralize SARS-CoV-2 yet potently neutralized pre-emergent and ancestral CoVs. One of these plasma-derived antibodies, SC45, exhibited *in vitro* potency against SARS-CoV-1 and the zoonotic “SARS-like” CoVs Pg-CoV, RsSHC014 and WIV1. Structural characterization of SC45 in complex with the Pg-CoV S2 domain revealed that SC45 engages a hairpin loop sandwiched between two N-linked glycosylation sites in the CD. Importantly, this epitope includes residues that are strongly conserved among clade 1a and 1b sarbecoviruses. To date, few antibodies have been reported to bind the CoV CD and only two antibodies – the murine mAb G4^45^ and human mAb F12^58^ which bind and neutralize MERS and HCoV-229E, respectively – have been structurally characterized. Notably, both G4 and F12 lack cross-reactivity to closely related CoVs within their respective genera (beta-CoV for G4 and alpha-CoV for F12) and neither neutralizes SARS-CoV-2. In contrast, SC45 binds to a distinct region upstream of both the G4 and F12 epitopes and displays high neutralization potency against zoonotic sarbecoviruses including Pg-CoV, WIV1, RsSHC014, and moderately neutralizes SARS-CoV-1. Furthermore, SC45 reduced weight loss and lung viral titers in mice infected with Pg-CoV. Of note it also conferred partial protection against SARS-CoV-2 challenge in mice, a finding which, in light of SC45’s lack of *in vitro* neutralization, suggests that some *in vivo* protection is likely mediated by Fc effector functions.

Multiple sequence alignment revealed that the SC45 epitope differs by a single amino acid substitution (D1084E) between SARS-CoV-2 and Pg-CoV, with an additional substitution (H1088Y) present in clade 1a coronaviruses. We observed that SC45 binds Pg-CoV and SARS-CoV-2 with comparable affinity, despite sequence variation at key contact residues. However, SC45 potently neutralizes WIV1 yet is approximately tenfold less potent against SARS-CoV-1, despite 100% sequence conservation of this epitope. Moreover, the substitution D1084E, which effectively recapitulates the Pg-CoV epitope in SARS-CoV-2, failed to sensitize the mutant SARS-CoV-2 virus to SC45-mediated neutralization. Our findings argue against sequence variability or binding affinity as primary determinants of neutralization and instead suggest that neutralization by SC45 may depend on disruption of long-range allosteric coupling between the CD epitope and conformational dynamics associated with membrane fusion^59–62^. However, further structural and functional studies will be necessary to define the precise molecular determinants of SC45’s neutralization activity. Although speculative, these data may suggest that SARS-CoV-2 evolved to escape neutralization from antibodies like SC45.

Structural and binding analyses show that SC45 is capable of engaging both the prefusion and postfusion conformations of the spike glycoprotein. This finding was unexpected given the structural reorganization of this domain during transitions from prefusion to postfusion. Notably, dual-prefusion and postfusion antibodies targeting “CD-like” membrane proximal epitopes have been described for class I fusion glycoproteins from Pneumoviridae^63–66^ and Paramyxoviridae^67,68^, where they neutralize viral entry by sterically blocking membrane fusion or by arresting refolding intermediates. Recent structural studies characterizing the refolding intermediates on the SARS-CoV-2 fusion pathway revealed that the CD is the principal “hinge” about which fusogenic rearrangements occur^69,70^. Thus, as an alternative hypothesis to allostery, SC45 binding to the CD may neutralize via mechanisms analogous to antibodies which engage basal epitopes on other class I viral fusion proteins.

Several S2-only vaccine constructs have demonstrated broad protective efficacy in animal models^39,71–74^. Whether antibody responses to the CD contributes meaningfully to protection against CoV infection remains unknown. Nonetheless, there is a growing appreciation that protection against SARS-CoV-2 can be mediated by non-neutralizing antibodies through Fc-dependent effector functions^75,76^, raising the possibility that CD-directed antibodies such as SC45 could contribute to protection *in vivo* despite limited neutralization, as we see in our data with partial protection (mitigated weight loss) against SARS-CoV-2.

Collectively, we define a new site of vulnerability in the CD of SARS-like viruses. This epitope exhibits three interesting features; (i) it is highly conserved among pre-emergent SARS-like viruses; (ii) it is sensitive to neutralization in multiple clade 1a and clade 1b sarbecoviruses, although notably not in SARS-CoV-2 suggesting that SARS-CoV-2 acquired distal mutations to escape neutralization and (iii) this epitope is accessible in both the prefusion and postfusion conformations. These findings expand the landscape of S2-directed epitopes in CoVs and highlight the potential for identifying neutralization-sensitive regions that can be targeted by broadly reactive S2 antibodies in human plasma. Finally, our data suggest that antibodies that moderately protect against circulating viral strains can recognize conserved and vulnerable epitopes on emerging CoVs. Despite the limited neutralizing potency of many S2-directed antibodies, cross-sarbecovirus binding is a recurrent feature in the serological repertoire and may provide a foundation for mitigating future zoonotic spillover events through Fc-mediated protection. Thus, vaccine strategies that focus on eliciting diverse antibody repertoires with complementary neutralization profiles may yield more robust and broad protection than strategies focused on eliciting highly potent antibodies at the expense of breadth.

Several limitations within our study warrant attention. First, although we demonstrate the protective efficacy of SC45 against Pg-CoV and partial protection against SARS-CoV-2 in mice, our study does not evaluate the impact of SC45 Fc-region modifications such as those which abrogate binding to Fc receptors. Additionally, we lack a high-resolution structure in the prefusion or with SARS-CoV-2 S2-antigen, each of which may provide additional mechanistic insight into SC45 neutralization. We also note that a K1086R mutation, which is located within the described CD epitope, emerged in multiple JN.1 subvariants^77^ and has since reverted to the ancestral 1086K through a recent recombination event^78^. Lastly, while we demonstrate broad neutralization by SC45, further discovery and characterization of a larger number of antibodies targeting the CD will be valuable for defining how vulnerabilities in the S2 and neutralization mechanisms vary across SARS-like CoVs and informing vaccine design strategies aimed at engaging highly conserved regions within the spike S2 subdomain.

## Methods

### Human subjects

The SARS-CoV-2 immune plasma and PBMC samples used in this study have been described previously^27^. Informed consent was obtained for all study participants under the University of Texas at Austin IRB protocol #2020-03-0085. In brief, whole blood was collected from four groups: (i) convalescent, non-hospitalized individuals with PCR-confirmed symptomatic SARS-CoV-2 infection; (ii) SARS-CoV-2–naïve individuals vaccinated with either the Pfizer BNT162b2 or Moderna mRNA-1273 vaccines; (iii) previously vaccinated but infection-naïve individuals who experienced a single breakthrough infection; and (iv) an individual who experienced two breakthrough infections. Blood was fractionated into plasma and PBMCs by density centrifugation using Histopaque-1077 (Sigma), and samples were frozen at −80 °C until analysis.

### Expression and purification of CoV spike proteins

Plasmids were transiently transfected in FreeStyle 293-F cells using polyethylenimine (PEI) and Kifunensine was added to a final concentration of 5 μM. Plasmids encoded the following: residues 1–1208 of the SARS-CoV-2 Wuhan S protein with a mutated S1/S2 cleavage site, proline substitutions at positions 817, 892, 899, 942, 986 and 987 and a C-terminal T4-fibritin trimerization motif, an 8×His tag and a TwinStrepTag (SARS-CoV-2 S6P); residues 1-1200 of the Pangolin coronavirus isolate Guangdong (Genbank ID OQ297708.1) with proline substitutions at positions 809, 884, 891, 934, 978 and 979, and a C-terminal T4-fibritin trimerization motif, an 8×His tag and a TwinStrepTag (Pg-CoV S6P); SARS-CoV-2 S6P with additional substitutions S383C-D985C to lock all receptor binding domain protomers in the down state (SARS-CoV-2 S6P-3D); residues 697-1208 with proline substituted at residues 817, 892, 899, 942, 986, 987, an engineered disulfide bond at S704C/K790C and an engineered salt-bridge Q957E, and C-terminal foldon trimerization motif of T4 fibritin, an HRV3C protease recognition site, an 8xHis tag, and a Strep-tag II (Hexapro-SS; SARS-CoV-2 S2-antigen); residues 689-1200 with proline residues 809, 884, 891, 934, 978 and 979, an engineered disulfide bond at S696C/K782C and engineered salt-bridge Q949E, and C-terminal foldon trimerization motif of T4 fibritin, an HRV3C protease recognition site, an 8xHis tag, and a Strep-tag II (Pg-CoV S2-antigen); residues 680-1191 of the CoV isolate WIV1 (Genbank ID KF367457.1) with profile substitutions at positions 800, 875, 882, 925, 969, 970, an engineered disulfide bond at S687C/K773C and an engineered salt-bridge Q940E, and C-terminal foldon trimerization motif of T4 fibritin, an HRV3C protease recognition site, an 8xHis tag, and a Strep-tag II (WIV1 S2-antigen). The supernatant was harvested 6 days after transfection, passed through a 0.22 uM filter and then applied to a StrepTactin column (IBA) for affinity purification. The 8xHis tag and Strep-tag II were cleaved with HRV3C protease at 4 °C overnight. Protein was further purified by size exclusion chromatography (SEC) using a Superose 6 10/300 column in SEC buffer (2 mM Tris pH 8.0, 200 mM NaCl, and 0.02% NaN3). For Ig-seq proteomic experiments, the SARS-CoV-2 S6P antigen was purified by SEC in PBS.

### VHonly repertoire sequencing

PBMCs were lysed in TRIzol Reagent (Invitrogen), and total RNA was extracted using the RNeasy Kit (Qiagen). First-strand cDNA was synthesized from 500 ng of RNA with SuperScript IV (Invitrogen). VH regions of the IgG, IgA, and IgM repertoires were then amplified from the cDNA using a multiplex primer set and the FastStart High Fidelity PCR System (Roche) under the following conditions: 2 min at 95°C; four cycles of 92°C for 30 s, 50°C for 30 s, 72°C for 1 min; four cycles of 92°C for 30 s, 55°C for 30 s, 72°C for 1 min; 22 cycles of 92°C for 30 s, 63°C for 30 s, 72°C for 1 min; 72°C for 7 min; hold at 4°C, as previously described. Products were concentrated separately in 15 uL of H2O using a PCR-purification kit (Zymo Research) and eluates were run on a 1% agarose gel to extract the full-length IgG, IgA and IgM amplicons. Following gel extraction and purification (Zymo Research), amplicons sequenced by 2×300 (bp) paired-end Illumina MiSeq.

### Native VHVL repertoire sequencing

Two million PBMCs from day 7 post-vaccination/infection were isolated into single-cell emulsion droplets containing magnetic oligo-d(T)25 magnetic beads (New England Biolabs) and lysis buffer (100 mM Tris pH 7.5, 500 mM LiCl, 10 mM EDTA, 1% lithium dodecyl sulfate, and 5 mM dithiothreitol) using a custom flow-focusing device as previously described. The magnetic beads were rescued from the oil emulsion by addition of hydrated ether. Beads were washed as previously described and resuspended in a RT-PCR solution with an overlap extension VH and VL primer set as previously described. Beads were re-emulsified using a DT20 dispersion tube (IKA) and aliquoted into a 96-well PCR plate at 100 uL per well. The overlap-extension RT-PCR was performed under the following conditions: 30 min at 55°C followed by 2 min at 94°C; four cycles of 94°C for 30 s, 50°C for 30 s, 72°C for 2 min; four cycles of 94°C for 30 s, 55°C for 30 s, 72°C for 2 min; 32 cycles of 94°C for 30 s, 60°C for 30 s, 72°C for 2 min; 72°C for 7 min; hold at 4°C. Magnetic beads were extracted from the emulsions with hydrated ether and pelleted on a magnetic rack. The supernatant containing the OE-RT-PCR products was concentrated using a PCR-purification kit (Zymo Research) and eluted in H2O. Amplicons were further amplified using a nested PCR, as previously described, and sequenced using 2×300 paired-end Illumina MiSeq.

### Proteomic analysis of anti-SARS-CoV-2 S6P human plasma IgG by mass spectrometry

Total IgG was purified from 1 mL of human plasma by affinity chromatography with Protein G Plus Agarose (Invitrogen), washed with 20 column volumes of PBS, eluted with 100 mM glycine-HCl pH 2.5, neutralized with 1M Tris-HCl pH 8.0 and buffer exchanged into PBS and concentrated using 10K MWCO Vivaspin centrifugal spin columns (Sartorius). Total IgG was cleaved into F(ab’)2 using IdeS. An affinity column was prepared as follows: 1 mg of SARS-CoV-2 S6P (1 mg/mL) was coupled to 50 mg of dry NHS-activated agarose resin (Thermo Fisher Scientific) overnight at 4C with rotation in PBS, remaining active sites were blocked with 2M ethanolamine for 20 minutes at room temperature with rotation, the column was washed 12x with 0.4 mL of PBS. Antigen specific IgG was isolated as follows: 3 mg of F(ab’)2 (10 mg/mL) was incubated with the SARS-CoV-2 6P resin for 1 hour at room temperature with rotation. Following F(ab’)2 binding, the resin was loaded into a spin column and centrifuged to collect the flow-through fraction. The column was washed 12x with 0.4 mL of PBS and eluted in 8 fractions with 0.5 mL of 1% formic acid. F(ab’)2 containing fractions were concentrated to dryness in a speed-vac, resuspended in LCMS grade H2O, combined and neutralized with 1M Tris/3M NaOH. The SARS-CoV-2 S6P-specific F(ab’)2 and 5 ug of the flow-through were separately prepared for bottom-up liquid chromatography-tandem mass spectrometry (LC-MS/MS) analysis as previously described. Samples were split into triplicates and loaded onto a C18 trap column for pre-concentration and desalting. Samples were then loaded onto a C18 analytical column and a gradient from 5 to 45% of mobile phase B (0.1% formic acid in acetonitrile) eluted peptides over 100 minutes with a total run time of 120 minutes. Survey scans were acquired at 120,000 resolution (at 200 m/z) over the scan range 400 – 1600 m/z in the Orbitrap, and the most abundant precursors were isolated in the quadrupole (1.6 m/z window) and fragmented using HCD (higher-energy collisional dissociation) within a 3 second cycle. Dynamic exclusion set to 45 seconds after two fragmentation events within a 30 second window. A targeted mass exclusion list containing m/z values for 88 constant region IgG peptides was also included. The resulting MS/MS fragment ions were detected in the linear ion trap.

### Bioinformatic analysis

Raw Illumina MiSeq output sequences were trimmed according to sequence quality using Trimmomatic version 0.27 and annotated using MiXCR version 2.1.6. Sequences with ≥ 2 full length VH reads were clustered into clonal lineages defined by 90% CDRH3 amino acid identity using USEARCH version 10.0.240. Donor specific MS search databases were constructed by concatenating bulk VHonly and natively paired VH and VL sequences from BCR-seq. MS/MS database searches were performed in Proteome Discoverer software version 1.4 (Thermo Fisher Scientific) with SEQUEST, as previously described. High-confidence peptide–spectrum matches (PSMs) from each precursor ion were retained for analysis. To ensure antigen specificity, only peptides with an elution-to-flowthrough extracted ion chromatogram (XIC) area ratio ≥ 5:1 were included after adjusting for differences in total XIC area between the elution and flow-through samples, as previously described. Relative abundance of SARS-CoV-2 S6 reactive plasma IgG lineages was then quantified by summing XIC areas of all VH peptides mapping to unique CDRH3s within each lineage and ranking lineages by total XIC area as a percentage of the total signal (plasma IgG repertoire). Longitudinal tracking of plasma lineages was performed by assessing clonal overlap across time points.

### Antibody expression and purification

Representative monoclonal antibodies (mAbs) from clonal lineages identified by Ig-seq and BCR-seq were synthesized as VH/VK/VL and cloned (Twist Biosciences) into their respective AbVec heavy (human IgG1 Fc constant) and light-chain expression vectors. Plasmids were mixed at a 1:3 molar ratio (VH:VL) and were transfected into Expi293F cells according to the manufacturer’s guidelines for Gibco Expi293 Expression System (Thermo Fisher Scientific). Cultures were incubated 37°C and 8% CO2 and harvested 5 days post-transfection. Antibody was purified by protein G plus agarose resin (Thermo Fisher Scientific) as described above and buffer exchanged into PBS with a10K MWCO Vivaspin centrifugal spin columns (Sartorius).

### Live virus neutralization assay

Live-virus neutralization assays were performed using full-length nLuc reporter virus constructs of SARS-CoV-2 D614G (Sequence Aq. No MT020880)^79^, Omicron BA.1 SARS-CoV-2 B.1.1.529 (EPI_ISL_6647961), Pangolin-CoV (EPI_ISL_410721), WIV1-CoV (KC881007.1), SARS-CoV 2003 (MK062183.1), SHC014-CoV (KC881005.1), were used, with the nano-luciferase (nLuc) reporter gene replacing CoV ORF 7a in SARS-CoV 2003, SARS-CoV-2, and Pangolin-CoV variants, and ORF 7 and 8 in WIV1 and SHC014-CoV, as previously published. Antibodies were prepared at 25 and 2.5 ug/mL and then serially diluted 3-fold on a 96-well plate for up to eight dilutions (Corning 3799) in virus growth medium (1X MEM [Gibco 11095080], 5% FBS [Hyclone SH30070.03HI] and 1% Penn-Strep [Gibco 10378016]). Plates were transferred into the BSL3 laboratory. Reporter virus individually diluted in the virus growth medium, added in equal volume too the mAb dilution plates, and incubated for 1 hour at 37C, 5% CO2. The virus+mAb dilutions were then transferred to duplicate columns on a 96-well black plate (Corning 3916) seeded with either Vero C1008 cells (ATCC CRL-1586) for sarbecoviruses, Huh 7.5 cells for MERS-CoV, and Huh7.5 cells stably expressing human IFITM3 for OC43-CoV, at 2 × 10E4 cells per well for a final virus dilution of 800 plaque-forming units (PFU) per well and incubated at 37°C, 5% CO2. After 18-24h incubation the virus growth for sarbeco- and merbecoviruses, and 48hr for OC43-CoV on each plate was quantified with the Promega Nano-Glo Luciferase Assay system (N1130) with a Promega GloMax Explorer (GM3500). The 50% inhibitory dilution (ID50) titer was defined as the serum dilution at which the observed relative light units (RLU) were reduced by 50% compared to virus+cell and virus-only control wells as determined by a Microsoft Excel macro and analyzed using GraphPad Prism 10.0.3. Additionally, all assays included monoclonal antibodies (ADG280 or S30942) serving as standardized assay performance controls.

### Generation of spike mutants

The molecular clone-derived SARS-CoV-2 WT and nLuc viruses were generated as previously described. To introduce the D1084E mutation into SARS-CoV-2, genomic cDNA sequences of SARS-CoV-2 was ligated *in vitro* and subjected to T7 transcription. The transcribed RNA was electroporated into Vero cells. SARS-CoV-2-ΔPRRA was generated by deleting the four residues PRRA coding sequence from the SARS-CoV-2 WT infectious cDNA clone^8^. All work was conducted in a high-containment BSL3 laboratory and personnel wore powered air purifying respirator (PAPR), Tyvek suits, aprons and booties, and were double gloved.

### Evaluation of mAb prophylactic activity in the K18-hACE2 mouse model

Mouse studies complied with all relevant ethical regulations and were performed after UNC IACUC protocol no. 20-074 review and approval. All infection studies were performed in animal biosafety level 3 (BSL-3) facilities at University of North Carolina at Chapel Hill. The hACE2-K18 mice (female 25 weeks old) used in these studies were obtained from the Jackson Laboratory (034860-B6.Cg-Tg(K18-ACE2)2Prlman/J). Mice (n=5 per group) were prophylactically treated with mAb SC45 at 200 ug intraperitoneally or with control isotype (Dengue virus mAb 2D22) 12 hours prior to infection, anaesthetized with a mixture of ketamine/xylazine and intranasally infected with 10^4^ plaque forming units of either Pangolin-Guangdong CoV MP789 SARS-CoV-2 Wuhan-Hu-1. Mice were monitored daily for clinical signs of disease (weight loss) and mortality. At indicated timepoints, mice were euthanized via isoflurane overdose and the lung tissue was harvested for viral lung titer analysis by plaque assay. After homogenization of lung tissue, the supernatant was serially diluted and used to infect 6-well plates containing monolayers of Vero E6 cells. Monolayers were overlayed with 0.8% agarose and plaques were visualized via red neutral dye 72 hours following infection.

### Determination of S2 subunit specificity by enzyme linked immunosorbent assay

Antibodies were screened by binding enzyme-linked immunosorbent assays (ELISAs) against in-house produced SARS-CoV-2 recombinant spike ectodomain proteins to identify S2 domain binding specificity. Indirect binding ELISAs were conducted in Costar 96-well plates (Corning) coated with either SARS-CoV-2 S6P or SARS-CoV-2 S2 at 4 ug/mL in phosphate buffered saline (PBS) overnight at 4C. Plates were blocked with PBS containing 2% milk (PBSM) for 2 hours at room temperature, washed 3x with PBS containing 0.1% (v/v) tween-20 and primary mAbs were incubated in PBSM at 15 ug/mL and serially diluted 3x from 15 ug/mL to 0.553 ug/mL. Plates were washed as described above and secondary antibody goat anti-human IgG (Fab)-horseradish peroxidase conjugate (Sigma-Aldrich) was diluted in PBSM 1:5000 and bound for 30 minutes at room temperature. Following washing, the secondary antibody was detected with 50 uL of 3,3’,5,5’-tetramethylbenzidine soluble substrate for 1 minutes and the reaction was quenched with 50 uL of 2M H2SO4. Absorbance at 450 nm was measured for each well using a Synergy H1 Microplate Reader (BioTek Instruments, Inc) and the area under the curve was determined from binding of serial dilutions.

### Surface plasmon resonance

Surface plasmon resonance (SPR) was performed by immobilizing prefusion-stabilized spike ectodomains (SARS-CoV-2 S6P, SARS-CoV-2 6P3D, Pg-CoV S6P, SARS-CoV-2 S2, Pg-CoV S2 and WIV1 S2) onto a Ni-NTA chip via its C-terminal His-tag. Two samples containing only running buffer (10 mM HEPES pH 8.0, 150 mM NaCl and 0.005% Tween 20) were injected over the ligand and reference flow cells, followed by injection of SC45 antigen-binding fragment (Fab) serially diluted from 400 – 3.125 nM. The chip was regenerated between cycles using 0.35 M EDTA and 0.1 M NaOH followed by 0.5 mM NiCl2. The resulting data were double-reference subtracted and fit to a 1:1 Langmuir binding model using the Biacore X100 Evaluation software.

### Cell-surface display of spike

Binding of SC45 to mammalian surface displayed spike was performed as previously described. SC45 was labeled with AF488 (Invitrogen) per the manufacturer’s instructions. On day 0, Expi293 cells (Thermo Fisher) were transfected with pEGFP alone (mock) or pEGFP with plasmid encoding either WT-SARS-CoV-2 Wuhan spike or WT-Pg-CoV spike in a 1:1 ratio in triplicate. After 24 hours, 3E5 cells were stained with a seven-point, fivefold serial dilution of SC45-AF568 beginning at 1000 nM, alongside an unstained control for 1 hour at 37C. Cells were washed with PBS containing 3% BSA and analyzed on a BD Fortessa flow cytometer. For each condition, data were obtained from two independent biological replicates, with two technical replicates per biological replicate. Data were processed in FlowJo by gating on forward and side scatter, singlets, and EGFP-positive events to identify successfully transfected cells. The geometric mean fluorescence intensity (GMFI) of the AF568 signal at each antibody concentration was used to determine the effective Kd as previously described.

### Cryo-EM sample preparation, data collection, and data processing

Glow-discharged UltrAuFoil 1.2/1.3 grids were prepared for 1 min at 0.39 mbar. The pangolin spike S2 subunit was incubated with a 1.5-fold molar excess of Fab per S2 monomer, yielding a final S2 trimer concentration of 0.6 mg/mL. The complex was incubated at room temperature for 5 min, after which amphipol was added to a final concentration of 0.1% (w/v). Aliquots were applied to grids and blotted for 4 s with a blot force of −3 prior to plunge-freezing. Cryo-EM data were collected on a 300 kV Titan Krios transmission electron microscope at a nominal magnification corresponding to a calibrated physical pixel size of 0.83 Å. Movies were recorded with a total exposure of 80 e⁻/Å². A total of 6,604 micrographs were collected, of which 5,127 were retained after quality control. From these, 1,317,793 particles were extracted, and 75,787 particles were selected for final refinement using cryoSPARC (v4.4.1)^80^. The final reconstruction was obtained at 2.91 Å resolution with imposed C3 symmetry, and the map was sharpened using DeepEMhancer^81^ via Cosmic2^82^. The postfusion SARS-CoV-2 spike protein structure (PDB ID 7BBH^83^) was used as the initial model and was iteratively rebuilt and refined using Phenix-1.21.2-5419^84,85^ Coot^86^ and UCSF ChimeraX ISOLDE^87,88^.

### Statistical analyses and software

Detailed information concerning the statistical methods used is provided in the figure legends. Flow data were analyzed using FlowJo version 10.5.3 (Treestar). Statistical analyses were performed with GraphPad Prism software version 8 (GraphPad). All cohorts started with five mice per harvest timepoint at the time of infection. Error bars indicate the SEM. We used Mann-Whitney U tests to compare two groups with non-normally distributed continuous variables. We used two-way analysis of variance (ANOVA) followed by Sidak’s multiple comparisons tests to analyze experiments with multiple groups and two independent variables. Experiments were performed independently at least twice to control for experimental variation. Comparisons are not statistically significant unless indicated.

## Supplemental Figures

**Figure S1.**
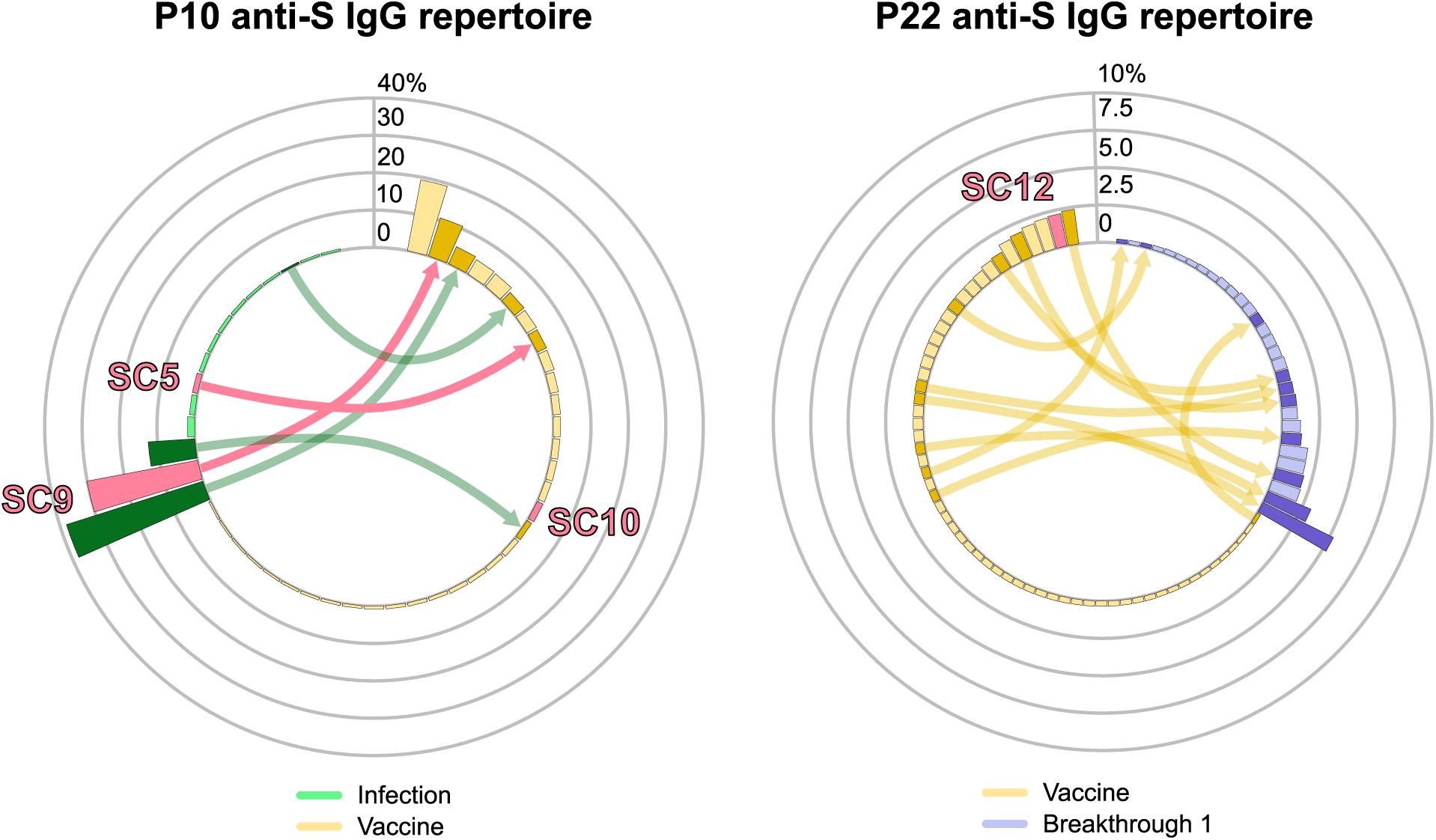
Plasma IgG repertoires across two additional donors with S2 mAbs indicated. Repertoire of SARS-CoV-2 spike-specific IgG in plasma in donor P10 (left), infected then vaccinated and P22 (right), vaccinated then infected. Only antibody lineages accounting for >0.5% of the plasma IgG repertoire are shown. Directional arrows connecting bars in the histogram demonstrate recalled lineages at each time point.

**Figure S2.**
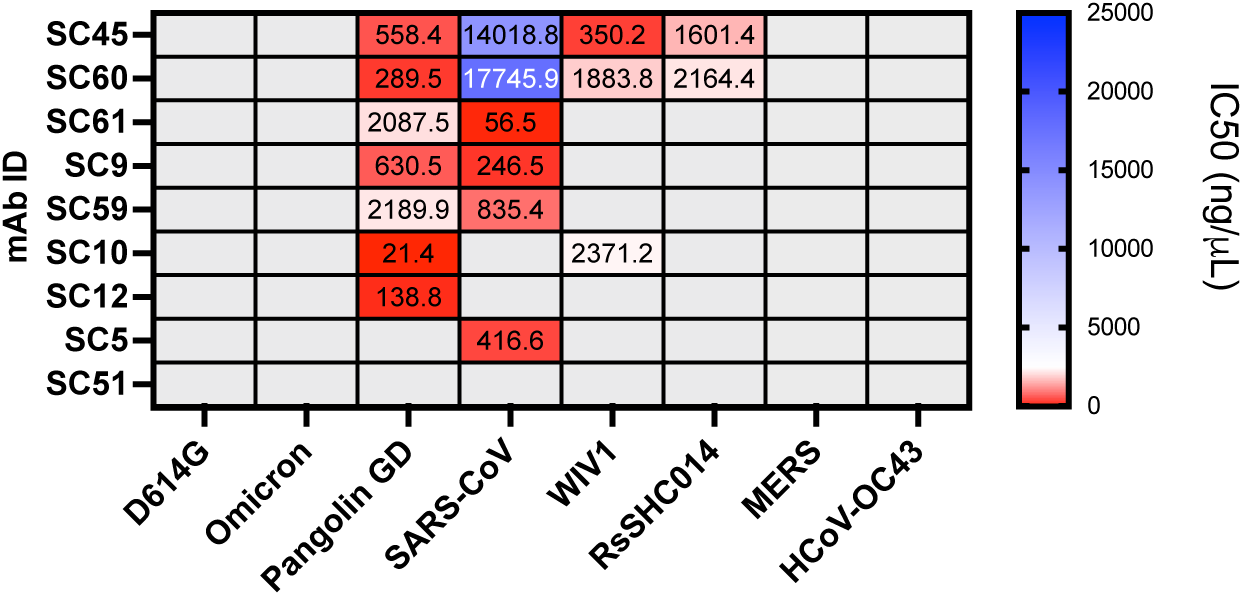
Live-virus neutralization assays using the S2-directed antibody panel with MERS-CoV and HCoV-OC43 viruses included. Non-neutralizing activity is shown in solid gray; related to Figure 1D.

**Figure S3.**
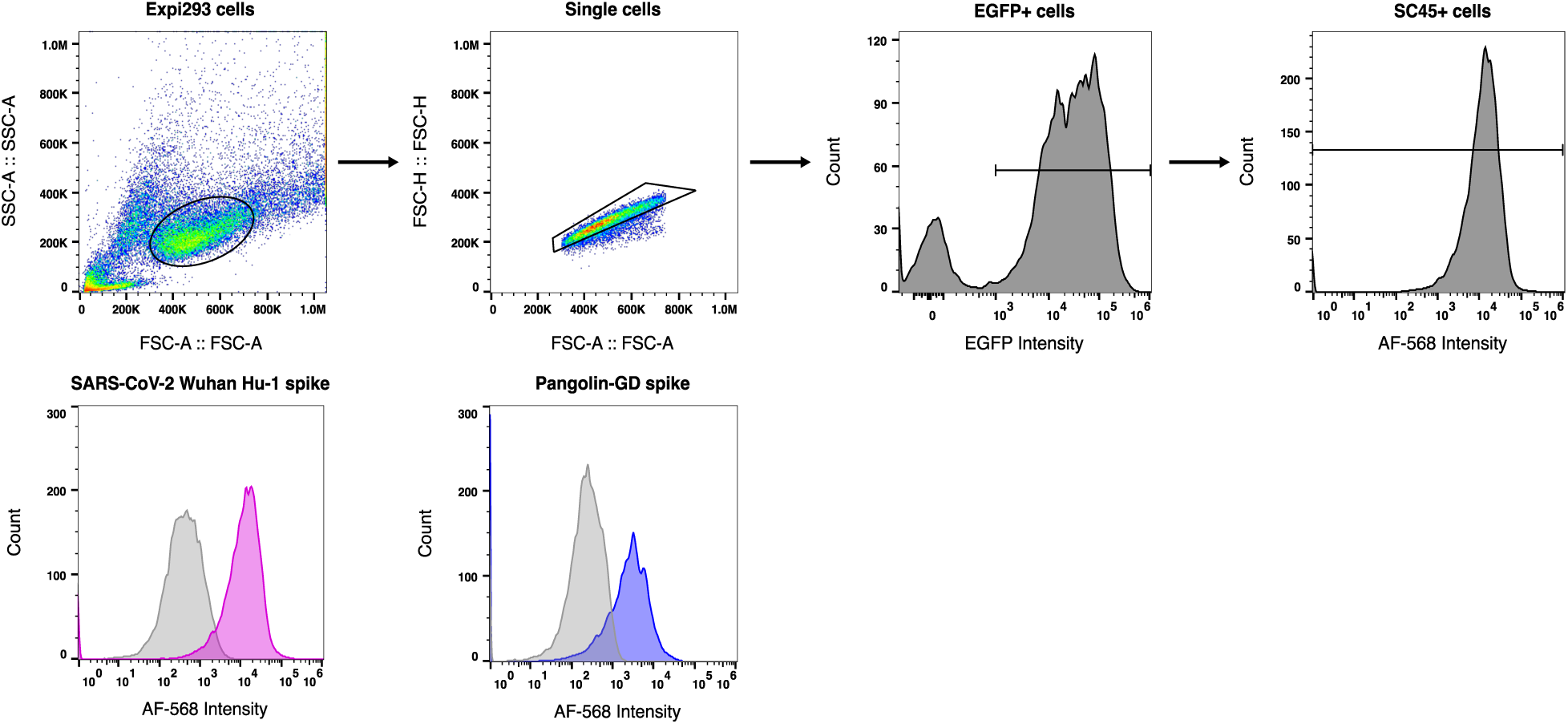
Gating strategy for analysis of SC45-AF568 binding to surface displayed spike on Expi293 cells. Representative plots demonstrating the sequential gating hierarchy used to isolate single Expi293 cells, identify successfully transfected EGFP+ populations, and quantify AF568 fluorescence (SC45 binding). This gating framework was uniformly applied across a titration gradient of SC45-AF568 to determine an effective antibody K_D_. Examples of AF568 positive SARS-CoV-2 Wuhan Hu-1 spike expressing and Pangolin GD spike expressing cells against respective mock transfected histograms after gating are shown.

**Figure S4.**
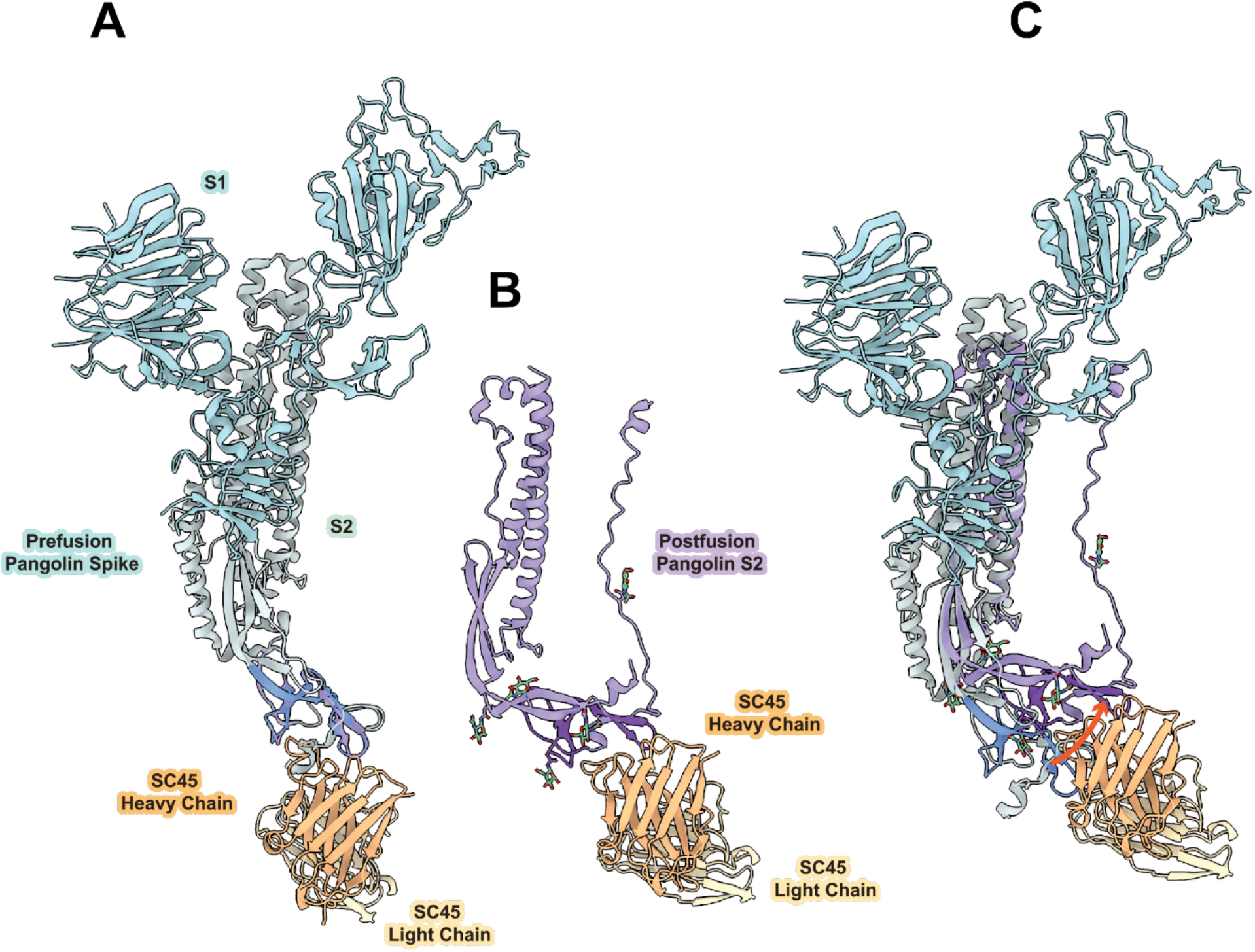
Comparison of prefusion and postfusion structures of Pg-CoV spike highlighting rigid-body translation of the connector domain. (A) Alignment of the solved Pg-CoV postfusion structure CD (purple) to the prefusion Pg-CoV spike S1 and S2 subunits (PDB ID: 7BBH). The prefusion connector domain (CD) is colored in cornflower blue. (B) Postfusion conformation of the solved Pg-CoV S2 subunit (residues 694-774, 982-1146), showing the SC45 antibody bound to a single protomer. (C) Overlay of the prefusion Pg-CoV structure with the SC45-Pg-CoV postfusion complex. The orange arrow indicates the rigid-body translation of the CD as the spike protein undergoes rearrangements from the prefusion-to-postfusion conformation.

**Figure S5.**
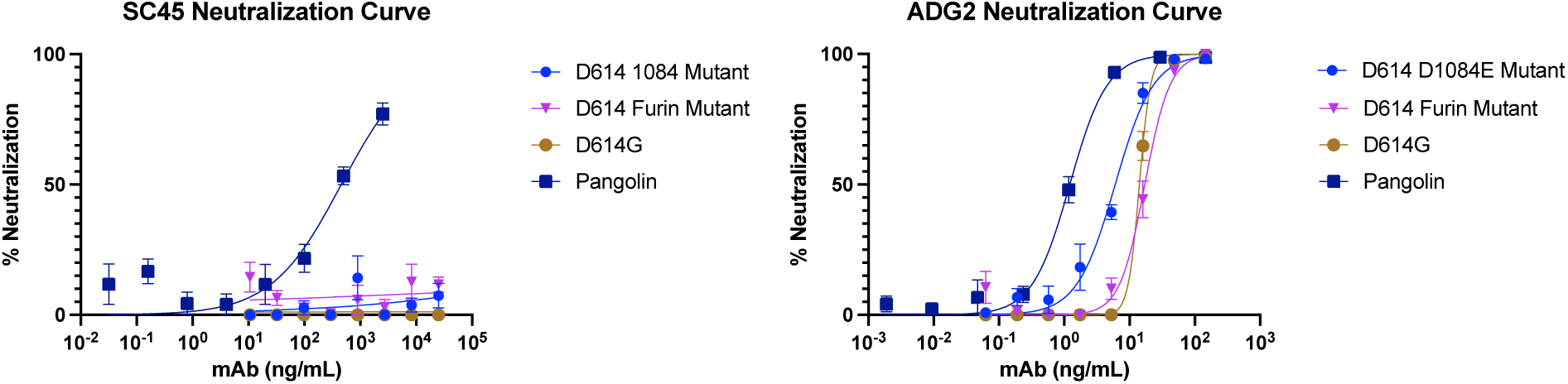
SARS-CoV-2 D614G furin knock-out and D1084E mutant viruses are resistant to SC45-mediated neutralization. Neutralization curves for SC45 (left) and positive control antibody ADG2 against Pg-CoV, SARS-CoV-2 D614G, SARS-CoV-2 D614G furin knock out mutant and SARS-CoV-2 D614G D1084E mutant. Results are shown from 2 independent experiments.

**Figure S6:**
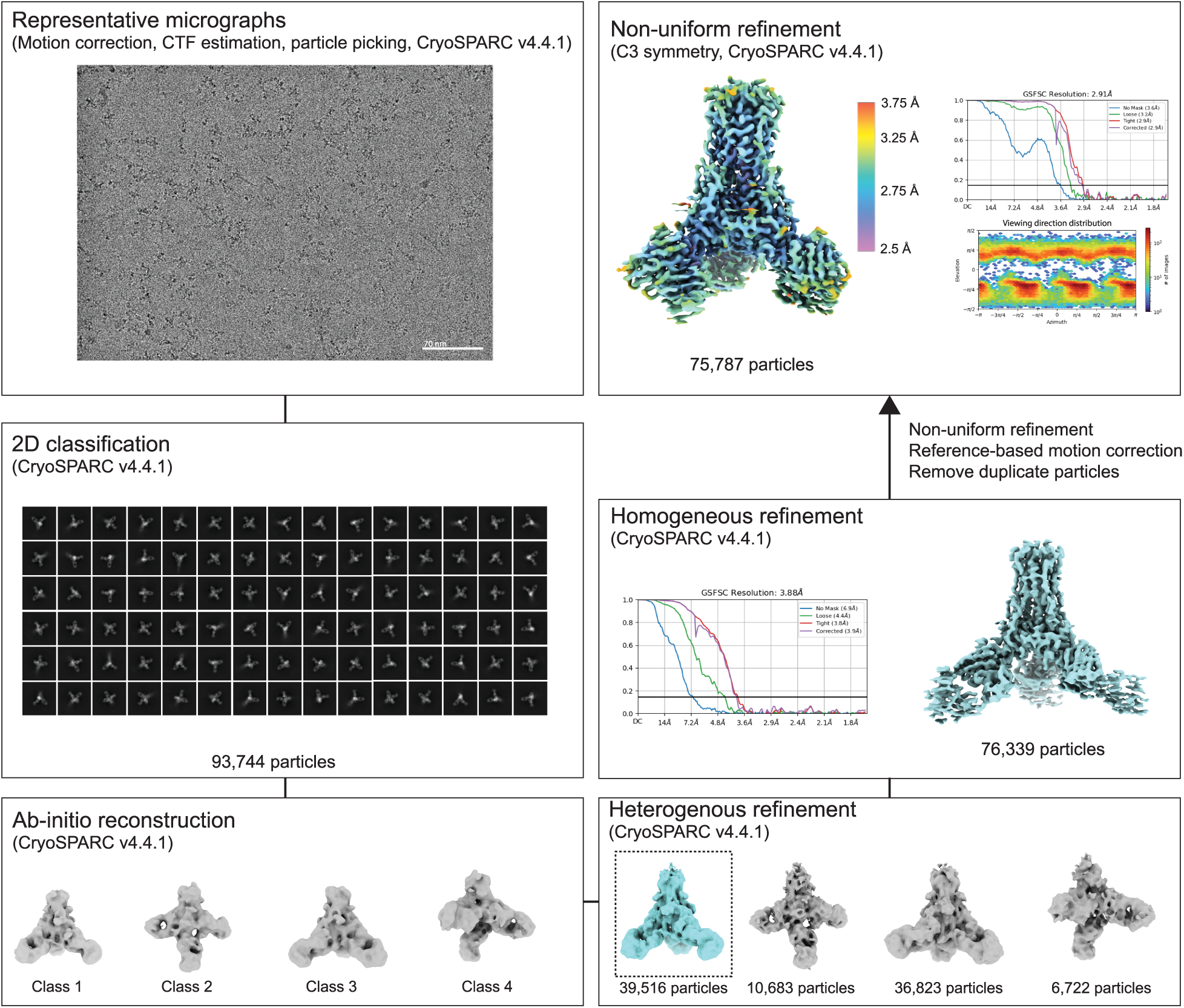
Cryo-EM processing summary for the Pg-CoV S2-Fab complex; related to Figure 4.

**Table S1:**
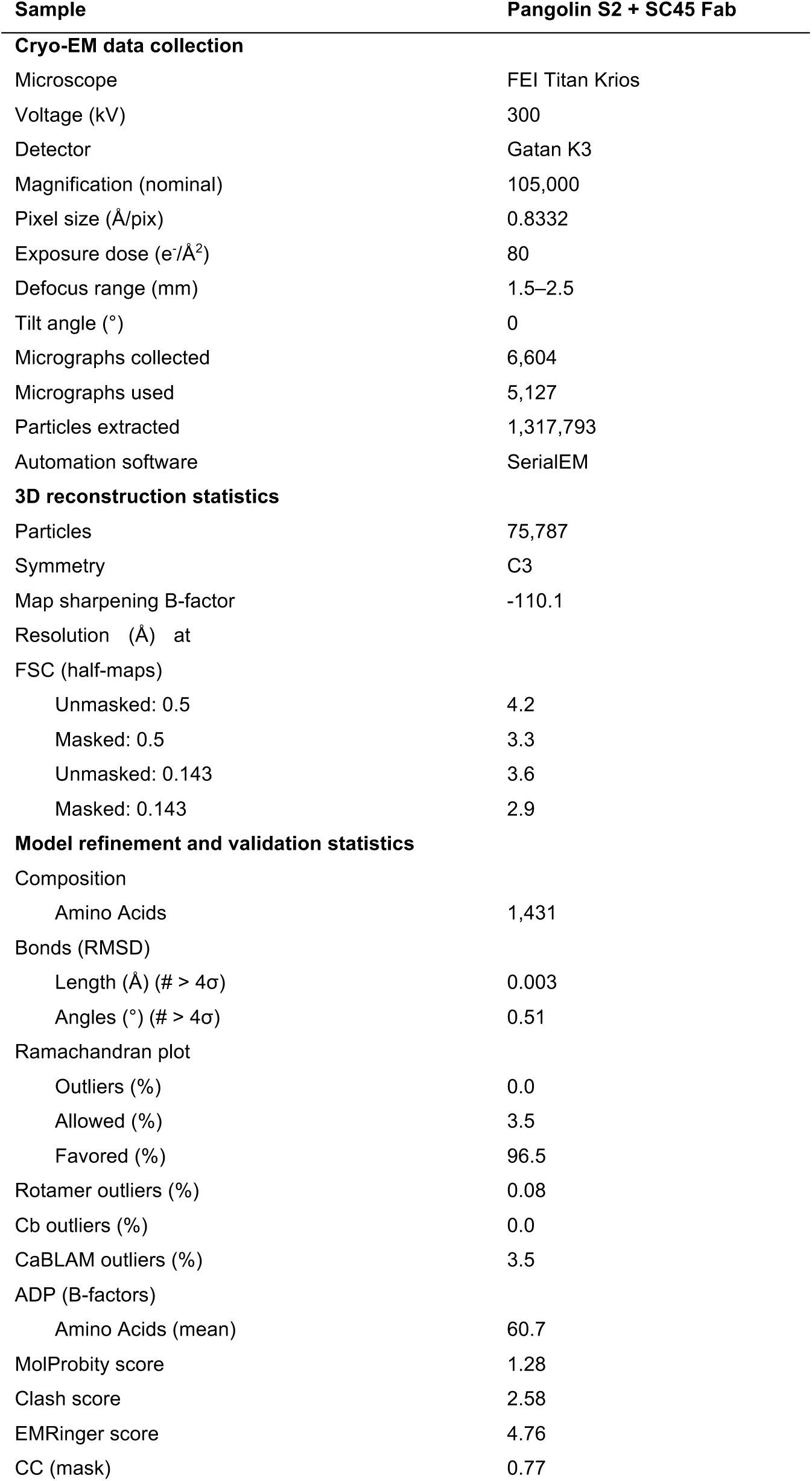
Cryo-EM data collection and refinement statistics; related to Figure 4.

| Sample | Pangolin S2 + SC45 Fab |
| --- | --- |
| **Cryo-EM data collection** |  |
| Microscope | FEI Titan Krios |
| Voltage (kV) | 300 |
| Detector | Gatan K3 |
| Magnification (nominal) | 105,000 |
| Pixel size (Å/pix) | 0.8332 |
| Exposure dose (e<sup>-</sup>/Å<sup>2</sup>) | 80 |
| Defocus range (mm) | 1.5–2.5 |
| Tilt angle (°) | 0 |
| Micrographs collected | 6,604 |
| Micrographs used | 5,127 |
| Particles extracted | 1,317,793 |
| Automation software | SerialEM |
| **3D reconstruction statistics** |  |
| Particles | 75,787 |
| Symmetry | C3 |
| Map sharpening B-factor | -110.1 |
| Resolution (Å) at FSC (half-maps) |  |
| Unmasked: 0.5 | 4.2 |
| Masked: 0.5 | 3.3 |
| Unmasked: 0.143 | 3.6 |
| Masked: 0.143 | 2.9 |
| **Model refinement and validation statistics** |  |
| Composition |  |
| Amino Acids | 1,431 |
| Bonds (RMSD) |  |
| Length (Å) (# > 4σ) | 0.003 |
| Angles (°) (# > 4σ) | 0.51 |
| Ramachandran plot |  |
| Outliers (%) | 0.0 |
| Allowed (%) | 3.5 |
| Favored (%) | 96.5 |
| Rotamer outliers (%) | 0.08 |
| Cb outliers (%) | 0.0 |
| CaBLAM outliers (%) | 3.5 |
| ADP (B-factors) |  |
| Amino Acids (mean) | 60.7 |
| MolProbity score | 1.28 |
| Clash score | 2.58 |
| EMRinger score | 4.76 |
| CC (mask) | 0.77 |

